# Decellularization enables functional analysis of ECM remodeling in planarian regeneration

**DOI:** 10.1101/2020.09.11.293936

**Authors:** Ekasit Sonpho, Frederick G. Mann, Michaella Levy, Eric J. Ross, Carlos Guerrero-Hernández, Laurence Florens, Anita Saraf, Viraj Doddihal, Puey Ounjai, Alejandro Sánchez Alvarado

## Abstract

The extracellular matrix (ECM) is a three-dimensional network of macromolecules that provides a microenvironment capable of supporting and regulating cell functions. However, only a few research organisms are available for the systematic dissection of the composition and functions of the ECM, particularly during regeneration. We utilized a free-living flatworm *Schmidtea mediterranea* to develop an integrative approach consisting of decellularization, proteomics, and RNA-interference (RNAi) to characterize and investigate ECM functions during tissue homeostasis and regeneration. High-quality ECM was isolated from planarians, and its matrisome profile was characterized by LC-MS/MS. The functions of identified ECM components were interrogated using RNAi. Using this approach, we discovered that heparan sulfate proteoglycan and kyphoscoliosis peptidase are essential for both tissue homeostasis and regeneration. Our strategy provides a robust experimental approach for identifying novel ECM components involved in regeneration that might not be discovered bioinformatically.

## Introduction

Tissue regeneration is an essential process for many organisms that can be activated during embryogenesis, throughout life during the constant physiological renewal of tissues, and in the dramatic restoration of missing body parts following injury or amputation (Galliot et al. 2017). The complexity of tissue regeneration is reflected by the numerous and overlapping molecular and cellular activities underpinning the restoration and integration of missing body parts (Poss 2007). In order for the organism to maintain and repair physiological form and function during and after tissue regeneration, cells need to communicate flawlessly with their microenvironment (Wang et al. 2013; Lukjanenko et al. 2016; Godwin et al. 2017).

The extracellular matrix (ECM), which is a collection of dynamically secreted and modified macromolecules occupying intercellular space, plays a central role in effecting cell-environmental communication (Daley, Peters, and Larsen 2008; Schultz and Wysocki 2009). Bidirectional crosstalk between cells and the ECM via secretion and selective degradation creates microenvironmental conditions capable of modulating cell proliferation, migration, differentiation and ultimately, the homeostatic maintenance of tissues throughout the lifetime of an organism (Koochekpour, Merzak, and Pilkington 1995; Bosnakovski et al. 2006; Zhen and Cao 2014; Bonnans, Chou, and Werb 2014). Of particular interest are the microenvironmental conditions governing stem cell biology (You et al. 2014; Morgner et al. 2015; Seyedhassantehrani et al. 2017).

Several animal models have been used for investigating how the ECM is involved in tissue regeneration processes, such as in *Hydra vulgaris* (Sarras 2012), axolotls (Phan et al. 2015), and zebrafish (Sánchez-Iranzo et al. 2018). Although progress has been made in these organisms implicating the ECM in regeneration, the inherent biology of these animals makes it challenging to systematically dissect ECM composition and to functionally study their possible roles in regeneration. For example, although *Hydra* is able to regenerate a whole animal from a clump of dissociated cells (Vogg, Galliot, and Tsiairis 2019) while axolotls (Phan et al. 2015) and zebrafish (Sánchez-Iranzo et al. 2018) can regenerate missing body parts, it is still difficult to carry out large-scale, loss-of-function screens in these adult organisms. Therefore, in an effort to systematically interrogate how ECM may contribute to whole-body and/or tissue regeneration, we chose to study the free-living freshwater planarian flatworm *Schmidtea mediterranea* (Sánchez Alvarado et al. 2002), which has extraordinary regenerative capacities and has been shown to be amenable to large-scale genetic interrogation (Reddien et al. 2005).

Because our current knowledge of ECM biology in planarians is limited (Isolani et al. 2013; Seebeck et al. 2017; Lindsay-Mosher, Chan, and Pearson 2020), it is first necessary to develop a comprehensive and optimized workflow to characterize and study the planarian ECM. A recent study has characterized the transcriptional landscape of ECM components in planarians by constructing an *in silico* matrisome (Cote, Simental, and Reddien 2019). However, this has not revealed the actual protein composition and distribution of these molecules. Similarly, Sonpho *et al*. successfully developed a simple technique for characterizing the morphology of isolated ECM by whole organism decellularization of a different planarian species. However; a complete systematic workflow for studying ECM biology in planarians has not yet been established (Sonpho et al. 2020).

Here, we propose an integrative workflow to systematically characterize the *S. mediterranea* ECM. This workflow consists of three core components: decellularization, proteomics, and RNA interference (RNAi) of identified ECM components. First, decellularization of *S. mediterranea* was optimized for the isolation of planarian ECM. Second, we subjected the decellularized fraction to biochemical characterization by liquid chromatography coupled to mass spectrometry (LC-MS/MS), which allowed us to construct the first experimentally derived planarian matrisome database. Third, proteins identified from proteomics allowed us to perform an RNAi screen to test their functions. In this work, we identified two ECM proteins that play important roles in both homeostasis and regeneration. In summary, by combining decellularization, proteomics, and RNAi screening, we provide proof-of-concept experimental evidence illustrating the potential of this workflow to discover and study ECM composition, function, and dynamics in an adult, regeneration-competent organism. Our approach also lays the foundation for a systematic, functional dissection of the role that the ECM may play in regulating stem cell behavior and function during both animal homeostasis and regeneration in planarians.

## Experimental Procedures

### Animal husbandry

*Schmidtea mediterranea* asexual clonal line CIW4 was maintained in 1X Montjuic salt solution, as described previously for static culture (Newmark and Sánchez Alvarado 2000). *S. mediterranea* were fed with beef liver once a week. The animals were starved for at least 1 week before experiments.

### ECM isolation

Whole-mount planarian decellularization has been optimized based on a previous publication (Sonpho et al. 2020). We optimized three protocols for decellularization, which we refer to as No Pretreatment ECM (NP-ECM), Formaldehyde ECM (FA-ECM), and N-Acetyl Cysteine pretreatment (NAC-ECM). All worms were collected from static culture. All three protocols were designed for 20 planarians.

For NP-ECM, planarians were incubated in 40 mL of 0.08% SDS decellularization solution for 18 hr at 4°C. ECM was harvested carefully using plastic transfer pipets and collected into a microtube. Decellularization solution was removed and replaced with 1 mL of wash solution for 30 min, 3 times, on ice. The ECM pellet was centrifuged at 14,000 *g* for 10 min.

For FA-ECM, planarians were fixed and stabilized by incubating in 0.8% formaldehyde solution at 4°C for 1 hr without shaking. After stabilization, the solution was replaced with 40 mL of 0.7% SDS decellularization solution for 18 hr at 4°C. To ensure complete decellularization, the solution was replaced with 40 mL 0.08% SDS decellularization solution for 2 hr at 4°C.

For NAC-ECM, planarians were first incubated with 5% N-Acetyl Cysteine (Sigma-Aldrich, MO) solution in 1X PBS for 10 min at RT on a seesaw rocker. Afterward, planarians were transferred into 1X PBS for a brief wash. Then, planarians were transferred into 40 mL of 0.08% SDS decellularization solution for 18 hr at 4°C. Subsequently, the solution was replaced with 40 mL of 0.04% SDS decellularization solution for 2 hr at 4°C. All of the solution components for *S. mediterranea* decellularization are reported in Table S1.

### Quantification of nucleotide contamination

Nucleic acid material remaining in ECM preparations was isolated using a DNeasy Blood & Tissue Kit (Qiagen, Germany). The amount of DNA was quantified using Qubit Fluorometric Quantification.

### Protein quantitation and Western blot analysis

Protein concentration for all samples was quantified by Pierce™ BCA Protein Assay Kit (Thermo Fisher, Waltham, MA). All samples were processed for analysis on SDS-PAGE. Briefly, ECM was harvested and solubilized by adding RIPA buffer and loading buffer (Biorad, Hercules, CA). Equal amounts of proteins were prepared and loaded per lane. PVDF membrane was activated by incubating in 100% Methanol for 5 min before washing in the transfer buffer. Proteins were transferred from the gel to the PVDF membrane with 100 V for 1 hr, then the PVDF membrane was washed with TBS-Tween 0.05% and blocked with 5% milk in TBS-Tween 0.05%, for 1 hr at room temperature (RT). A dilution of primary antibody was prepared in a fresh blocking solution as follows: rabbit histone H3 antibody diluted at 1:4000, and rabbit MMP3 antibody diluted at 1:1000. The blocking solution was replaced with the primary antibody solution overnight at RT. Membrane was then washed with TBS-tween 0.05%, 4 times for 15 min each. The HRP-secondary antibody against rabbit antibody was diluted at 1:5000 in 1% milk. The membrane was incubated with the secondary antibody for 1 hr at RT. The membrane was washed with TBS-tween 0.05%, 4 times for 15 min each. The signal of HRP was visualized via SuperSignal™ West Femto Maximum Sensitivity Substrate (Thermo Fisher, Waltham, MA). The membrane was developed until the signal appeared. Exposures for Histone H3 and MMP3 antibodies were 1 min and 2 hr, respectively.

### Histological staining

Worms were processed using a Delta Pathos hybrid tissue processor (Milestone Medical). Briefly, after fixing planarians in 1% Nitric acid, 50 μM MgSO4 and 0.8% formaldehyde overnight, tissues were rinsed and dehydrated in 70% ethanol. The following steps were automatically performed by the Delta Pathos processor: 4 min rinse with 70% ethanol; 100% ethanol for 10 min at 65°C; 100% isopropanol for 30 min at 68 °C. For paraffin infiltration, the first bath of paraffin was set at 70°C for 8 min, then, fresh paraffin was replaced for an additional 6 min, followed by a final step of fresh paraffin incubation for another 20 min at 65°C (Tissue Infiltration Medium by Surgipath available through Leica). The paraffin embedded tissue was sectioned in Richard-Allan Type 9 paraffin (Thermo Scientific). Tissue was sectioned at 5 μm thickness, dried, heated, then stained with Masson’s Trichrome Stain. This procedure is derived from *Histotechnology, a Self-Instructional Text* (Carson and Hladik 2009).

### Scanning Electron Microscopy

To prepare samples for observation under a scanning electron microscope (SEM), collected planarian ECM was incubated in fixative solution composed of 2.5% glutaraldehyde, 2% paraformaldehyde, 1mM calcium chloride, 1% sucrose in 50mM sodium cacodylate buffer. Afterward, the sample was stained with tannic acid, osmium, and thiocarbohydrazide, as previously described (Jongebloed et al. 1999). At the end of dehydration, the samples were critical point dried in a Tousimis Samdri 795 critical point dryer, then mounted on stubs and coated with 5 nm of gold palladium in a Leica ACE600 sputter coater. The samples were imaged using a Zeiss Merlin SEM at 20 kV and 400 pA.

### Proteomics analysis

All decellularized samples were centrifuged at 16,900 *g* at 4°C to recover the ECM pellets. Control samples consisting of whole animal protein extracts were also prepared for comparison by freezing whole planarians in liquid nitrogen. ECM and control pellets were resuspended in 120 μL of 100 mM Tris-HCl, pH 8.5, and 8 M Urea. The samples were vortexed vigorously to dissolve pellets. To reduce disulfide bonds, Tris(2-carboxyethyl)phosphine hydrochloride (TCEP, Pierce) was added to 5 mM (0.6 μL of 1 M of TCEP) and incubated at RT for 30 min. Reduced cysteines were alkylated by adding 2.4 μL of a freshly made 0.5 M chloroacetamide stock solution (CAM, Sigma) and incubating at RT in the dark for 30 min. Samples were deglycosylated with PNGase F by adding 5μl of a solution consisting of 18μL of Glycobuffer 2 (10x) and 27μL of PNGaseF (NEB, P0708S). Endoproteinase Lys-C (Promega), 5 μL at 1 μg/μL, was added and samples were incubated at 37 °C overnight while shaking. Solutions were diluted to 2 M urea by adding 360 μL of 100 mM Tris-HCl and 2 μL of CaCl_2_. For trypsin digestion, 10 μL of Trypsin (Promega) at 0.1 μg/μL was added and samples were incubated at 37 °C overnight while shaking. Reactions were quenched by adding 90% formic acid to a final concentration of 5%. After digestion, peptide concentrations were quantitated by the Pierce Quantitative Colorimetric Peptide Assay (Thermo Fisher).

Peptides were diluted with Buffer A (5% acetonitrile (ACN), 0.1% formic acid (FA)). Then, samples were injected on an in-house packed 50 μm inner diameter microcapillary column packed with 10 cm of 1.9 μm Aqua C18 resin (Dr. Maisch GmbH). The samples were injected in 100% Buffer A and subsequently eluted using an Ultimate 3000 UPLC (Dionex) at a flow rate of 0.120 μL/min for 240 min from 10-40% Buffer B (80% ACN, 0.1% FA) before ramping to 95% B in 25 min. The high organic concentration was maintained for 15 min before re-equilibration and the reinjection with the next sample. Peptides were ionized by the application of a 2.5 kV positive voltage applied distally to the peptides eluting into a Q-Exactive Plus (QE+) mass spectrometer (Thermo Scientific, Waltham, MA). Full MS spectra were recorded on the eluting peptides over a 350 to 1700 m/z range at 70,000 resolving power. The AGC (automated gain control) target was 1×10^6^ with a max injection time set to 50 ms. The top 15 peptides with charges 2-5 were fragmentated by High energy Collision Dissociation (HCD) at 27% normalized collision energy (NCE) and the MS/MS fragmentation was collected at 17,500 resolving power, with an AGC target of 1×10^5^ and a maximum injection time set to 150 ms. Dynamic exclusion was enabled for 30 seconds (Zhang et al. 2009). Mass spectrometer scan functions and HPLC solvent gradients were controlled by the XCalibur data system (Thermo Scientific, Waltham, MA).

RAW files were extracted to .ms2 file format (McDonald et al. 2004) using RawDistiller v.1.0 (Zhang et al. 2011). RawDistiller settings were used to abstract MS1 scan profiles by Gaussian fitting and to implement dynamic offline lock mass using six background polydimethylcyclosiloxane ions as internal calibrants (Zhang et al. 2011). MS/MS spectra were first searched using ProLuCID v. 1.3.3 (Xu et al. 2015) with a peptide mass tolerance of 10 ppm and 25 ppm for peptide and fragment ions, respectively. Trypsin specificity was imposed on both ends of candidate peptides during the search against a protein database containing 30,863 *S. mediterranea* predicted protein sequences as well as 483 usual contaminants such as human keratins, IgGs and proteolytic enzymes. To estimate false discovery rates (FDR), each protein sequence was randomized (keeping the same amino acid composition and length) and the resulting “shuffled” sequences were added to the database, for a total search space of 62,564 amino acid sequences. A mass shift of 57.0215 Da was statically added to cysteine residues to account for alkylation by CAM, mass shifts of 15.9949 and 0.98401 Da were differentially added to methionine and asparagine residues, respectively, to account for oxidation due to sample handling and deglycosylation by PNGaseF.

DTASelect v.1.9 (Tabb, McDonald, and Yates 2002) was used in combination with an in-house software, swallow v1.0 (https://github.com/tzw-wen/kite), to select and sort peptide/spectrum matches (PSMs) to FDRs of less than 1% at the peptide and protein levels. Results from the worm without decellularization (3 replicates of control), formaldehyde-treated (3 replicates of FA-ECM), N-acetyl cysteine-treated (3 replicates of NAC-ECM), and not pretreated (2 replicates of NP-ECM) ECMs were merged and compared using CONTRAST v 1.9 (Tabb, McDonald, and Yates 2002) in combination with sandmartin v1.0, an in-house software, to select proteins at FDRs less than 5% when combining all runs. Proteins that were subsets of others were removed using the parsimony option in DTASelect on the proteins detected after merging all runs. Proteins that were identified by the same set of peptides (including at least one peptide unique to such protein group to distinguish between isoforms) were grouped together, and one accession number was arbitrarily considered as representative of each protein group.

NSAF7 v. 0.0.1 (Zhang et al. 2010) was used to create the final reports on all detected peptides and non-redundant proteins identified across the different runs and calculate label-free dNSAF quantitative values for all detected protein/protein groups. Spectral and protein level FDRs were, on average, 0.34 ± 0.10% and 1.5 ± 0.5%, respectively. The mass spectrometry datasets have been deposited to the ProteomeXchange Consortium via the PRIDE partner repository with the dataset identifier PXD013181 and 10.6019/PXD013181 (Deutsch et al. 2017; Perez-Riverol et al. 2019).

### Statistical analysis of differential protein expression

QPROT (Choi et al. 2015) was used to calculate log fold changes and false discovery rates (FDR) to identify proteins enriched in replicate analyses of each of the three ECM preparations, compared to the controls. Volcano significance plots were generated in Microsoft Excel by plotting the log fold change (X-axis) and the −log_10_(FDR) from QPROT (Choi et al. 2015) of each protein from the three types of ECM preparations compared to the control. Significance cut-offs were log_2_ fold change >2 and FDR <0.05. Dashed lines were added to the graphs at log_2_ fold change = 2 and −log_10_(FDR) = 1.3 (−log_10_(0.05)). Proteins in the upper right quadrant were identified as significantly enriched in ECM samples compared to controls. The full list of proteins identified in each condition is provided in Table S2.

### Gene ontology (GO) enrichment

*S. mediterranea* gene models were assigned Gene Ontology (Michael et al. 2000) terms by combining the GO annotations from PLANMINE (Rozanski et al. 2019), BLAST2GO (version: 5.2) (Götz et al. 2008), AHRD (version: 3.3.3), Uniprot/SWISS-PROT best BLAST hits (param: -evalue=.001; db date: 20180501) (Bateman 2019), InterProScan (version: 5.32-71.0) (Jones et al. 2014) and PANTHER. (Thomas et al. 2003). GO enrichment was performed using TopGO (version: 2.34.0) (Alexa and Rahnenfuhrer 2018).

### RNAi food preparation and feeding

The RNAi food was prepared as previously described (Rouhana et al. 2013). The primers used for cloning each gene are shown in Table S3. The colored liver puree was added to the bacteria pellet and fed to planarians cultured in an automatic water exchanger system (Arnold et al. 2016). Ten feeding cycles were implemented in both primary screening and secondary screening prior to use in following experiments. The essential transcription factor *nkx-2.2* was used as a positive control (Forsthoefel et al. 2012) while the *C. elegans* gene *unc-22* was used as a negative control for RNAi treatment (Reddien et al. 2005).

### Homeostasis and regeneration assays

Phenotypes of the targeted ECM components in tissue homeostasis were evaluated by observing unamputated planarians at 2 days after completion of the RNAi feeding. Amputations were performed by arranging planarians on a wet filter paper placed on a cold aluminum plate buried in ice and cutting with a surgical blade. The surgical blade was cleaned with 70% ethanol between amputations. Any signs of abnormal body plan or lysis were considered phenotypes. Worms were evaluated 10 days after sagittal amputation for morphological regeneration phenotypes. Any signs of incomplete or lack of newly regenerated tissue (blastema) were considered a defective phenotype. Worms that showed no touch response were considered dead.

### Whole-mount in situ hybridization (WISH) and animal radiation

A GammaCell 40 Exactor irradiator was used to expose planarians to 6,000 rad for lethal irradiations. For the WISH experiment, the protocol was as previously described (Pearson et al. 2009).

### Determination of cell-type specific expression of genes of interest

To access the published single cell RNA-seq database, we used the Planaria SCS online tool Digiworm (Fincher et al. 2018). The partial or complete contig of gene IDs was entered into the search box. For matching genes to the planarian transcriptome, the transcriptome assembly was mapped to the dd_v4 transcriptome as previously indicated (Brandl et al. 2016). The transcriptome assembly ID (dd_v4) was entered into the search box to generate Seurat maps.

### Statistical analysis of data

Statistical analysis was carried out by using the R software version 3.14 (RS team, 2018). A Two-way ANOVA was performed for control and treatment in all experiments. The protected LSD *post hoc* test (a=0.05) was used for multiple comparison tests between control and treatment. The hypergeometric test was used for the calculation of the statistics from Venn Diagrams.

## Results

### High-quality ECM can be successfully isolated from S. mediterranea

A planarian decellularization procedure used for removing cells from whole body planarians that leaves behind an extracellular matrix structure in the original shape of the animal has been recently described for *Dugesia japonica* (Sonpho et al. 2020). Since this protocol has not been tested for the isolation of ECM from *S. mediterranea*, we explored detergent-based tissue decellularization at low temperature for isolating extracellular matrix in *S. mediterranea* by incubating planarians with a low concentration of SDS for 18 hours at 4 °C. Compared to our intact control animals (Figure 1A), the results showed that decellularization without any pretreatment (NP-ECM) could maintain the animal’s overall morphology, yet, the external structure of the ECM was tremendously compromised and the ECM’s internal structures were not well-defined (Figure 1B). In order to preserve external and internal integrity of isolated ECM, formaldehyde was added to stabilize the ECM structure prior to the decellularization (Sonpho et al. 2020). As expected, formaldehyde stabilization yielded a translucent ECM (FA-ECM) with intact gross external structures and maintained the organization of internal organs such as the digestive tract branches, eye spots, and pharynx (Figure 1C).

**Figure 1.**
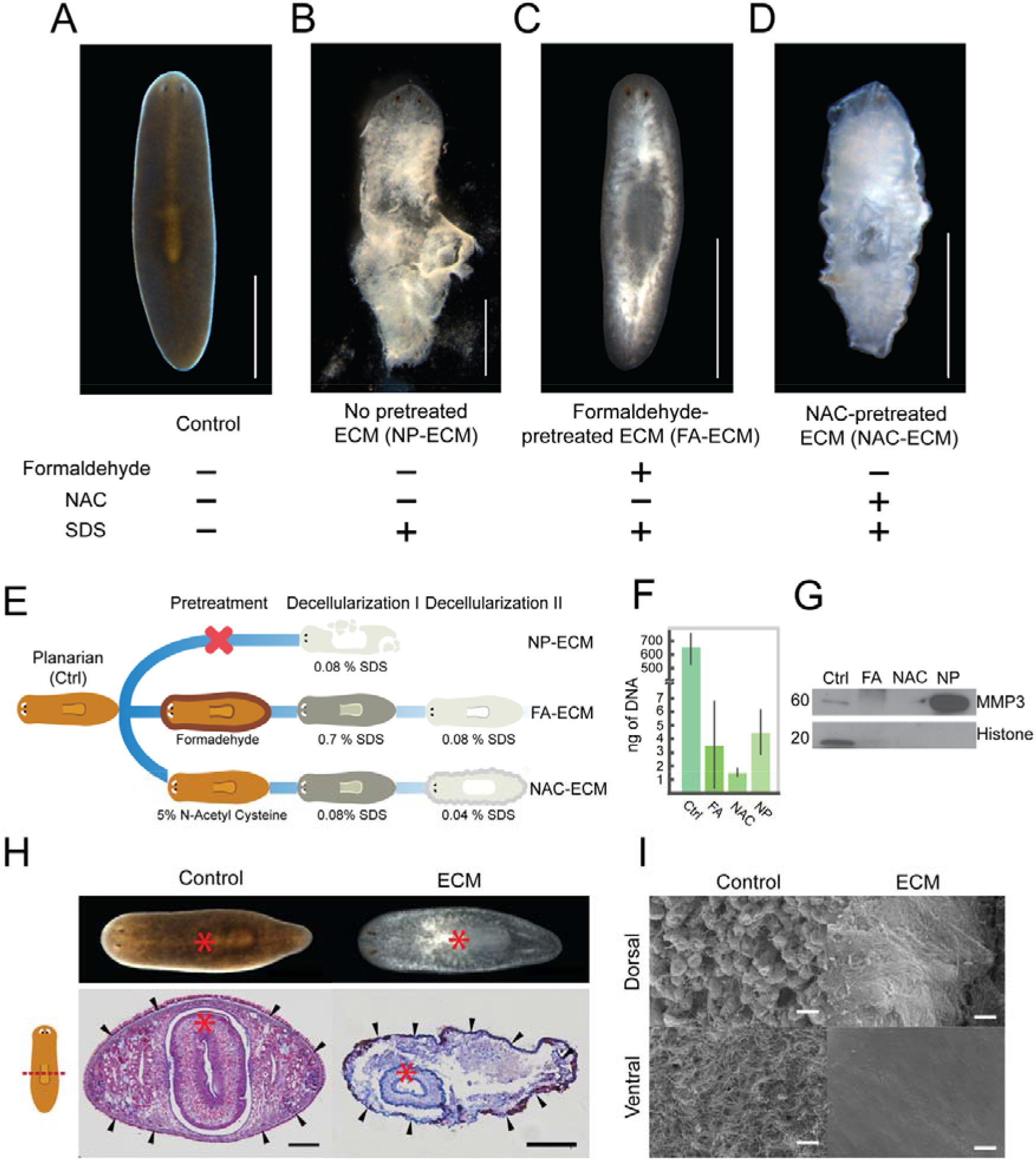
Isolation of *S. mediterranea* ECM by tissue decellularization. Overall morphology of (A) a control worm (Ctrl) and ECMs isolated from different pretreatment procedures, including (B) no pretreatment (NP-ECM), (C) formaldehyde-treated ECM (FA-ECM), and (D) N-Acetyl cysteine-treated ECM (NAC-ECM). Scale bars represent 1 mm (A-D). (E) The detailed protocol for ECM isolation. (F) Amount of total DNA removed by the decellularization procedure. Error bars represent standard deviation. The data were calculated from 3 independent experiments, ANOVA, (p < 0.05). (G) Western blot against MMP3 and Histone H3. Equal amount of total protein was loaded into each lane. (H) Gross morphology and histological sections of decellularized planarians stained with Masson’s trichrome. Decellularizing FA-ECMs were collected at 18 hr. Black arrows indicate layer of basement membrane. Red asterisk indicates the location of pharynx. Scale bars represent 500 μm. (I) The external surface topology of planarians and FA-ECM were visualized by scanning electron microscopy. Scale bars represent 10 μm.

This result suggests that chemical pretreatment of animals prior to decellularization could be critical for preserving intact structures of the decellularized organism. In order to support this hypothesis, N-acetyl cysteine (NAC) was used to pretreat animals prior to decellularization. Our result indicates that NAC pretreatment prior to decellularization (NAC-ECM) could also preserve decellularized ECM but to a lesser extent than FA-ECM (Figure 1D). Therefore, we concluded that pretreatment of animals prior to whole body decellularization is imperative for successfully isolating structural intact ECM from whole planarians. Additionally, pretreatment of animals via formaldehyde fixation yields an external structure that is better preserved than either NAC-ECM or NP-ECM. Furthermore, the decellularization process was also captured by time-lapse movies for providing the real-time dynamics of whole animal decellularization (Video S1). The footage displayed the reduction of animal size during tissue decellularization in all three decellularization protocols. There was a small change in size from FA-ECM whereas there was a dramatic change in NAC-ECM. Meanwhile, NP-ECM displayed a thoroughly compromised structure as well as the disruption of the basement membrane.

In summary, we successfully implemented three approaches for decellularizing whole *S. mediterranea*. The detailed protocols for these three ECM isolation approaches are summarized in Figure 1E.

### Extracted ECM is free of cells

Although decellularization yielded a promising ECM structure based on the visual inspection of whole planarians, intact cells may have remained in the interior of the planarian body. Therefore, cellular contamination in all three ECM extraction protocols was evaluated by measuring the quantity of DNA. The results indicated that the amount of DNA per worm in the three ECM preparations was at least two orders of magnitude lower when compared to controls (Figure 1F). The absence of cells was further validated by western blot analysis. An antibody against histone H3 was used as the marker for intracellular proteins and metalloproteinase-3 (MMP3) was used as the marker of ECM. The results indicated that histone H3 was detected only in the control whilst FA-ECM showed a very faint band of histone H3 (Figure 1G). NAC-ECM and NP-ECM showed no visible band of histone H3. Additionally, the antibody against MMP3 revealed the presence of MMP3 in both control and NP-ECM. The intensity of the MMP3 band in the control was much fainter than the analogous band in NP-ECM, suggesting an enrichment of the ECM fractions by decellularization. Taken together, the absence of detectable DNA and histone H3 suggested that the three decellularization protocols are able to isolate ECM that is nearly free of cells.

### Structures are maintained in extracted ECM

As an additional means of testing for the presence of expected ECM structures, we performed two morphological characterizations on the most structurally preserved ECM, FA-ECM. First, the preservation of the basement membrane post-decellularization was assessed by Masson’s trichrome staining. The result demonstrated that decellularization was able to infiltrate epithelial cells and remove mesenchymal cells based on the depletion of pink and purple in ECM samples as compared to the control whilst maintaining the integrity of the basement membrane, indicated in blue at the outermost region of ECM labelled with black arrows (Figure 1H). The results also confirmed that internal structures are preserved during decellularization, as evidenced by the intact pharynx. Second, the external surface of planarian ECM was examined by scanning electron microscopy (SEM). The dorsal and ventral sides of control animals were covered with a mucus layer and a ciliated surface, respectively (Figure 1I). However, for FA-ECM, a fibrous surface was observed on the dorsal side while a non-ciliated surface was observed ventrally. This suggests that the decellularization process removes the mucus layer and epithelial cells as well as the ventral ciliated surface. On the whole, our two morphological characterizations confirm the cell-free environment of our FA-ECM as well as the preservation of ECM integrity with an intact basement membrane.

### Proteomics analysis of decellularized planarians reveals complex matrisome landscape

The decellularization protocol provided us with an ECM enriched sample allowing us to empirically determine the protein composition of ECM (Figure 1G). We expected ECM proteomic profiles to differ from control as decellularization ought to remove intracellular proteins and non-ECM extracellular proteins. We further assayed the reduced complexity of the ECM samples through SDS-PAGE. The differential pattern of protein bands of SDS-PAGE between ECM and control samples supports the hypothesis that intracellular proteins have been depleted by decellularization (Figure S1). We used mass spectrometry to determine the identity of the proteins in the ECM via the workflow illustrated in Figure 2A. The number of non-redundant (NR) spectra, NR peptides, and NR protein counts obtained from the four types of samples are shown in Figure 2B and Table S4. These results show that all ECM samples have fewer NR spectra, NR peptides, and NR proteins compared to the whole animal control, suggesting that intracellular proteins were depleted by decellularization. Amongst all ECM samples, NAC-ECM shows the highest NR spectra, NR peptide, and NR protein counts while FA-ECM displays the lowest counts.

**Figure 2.**
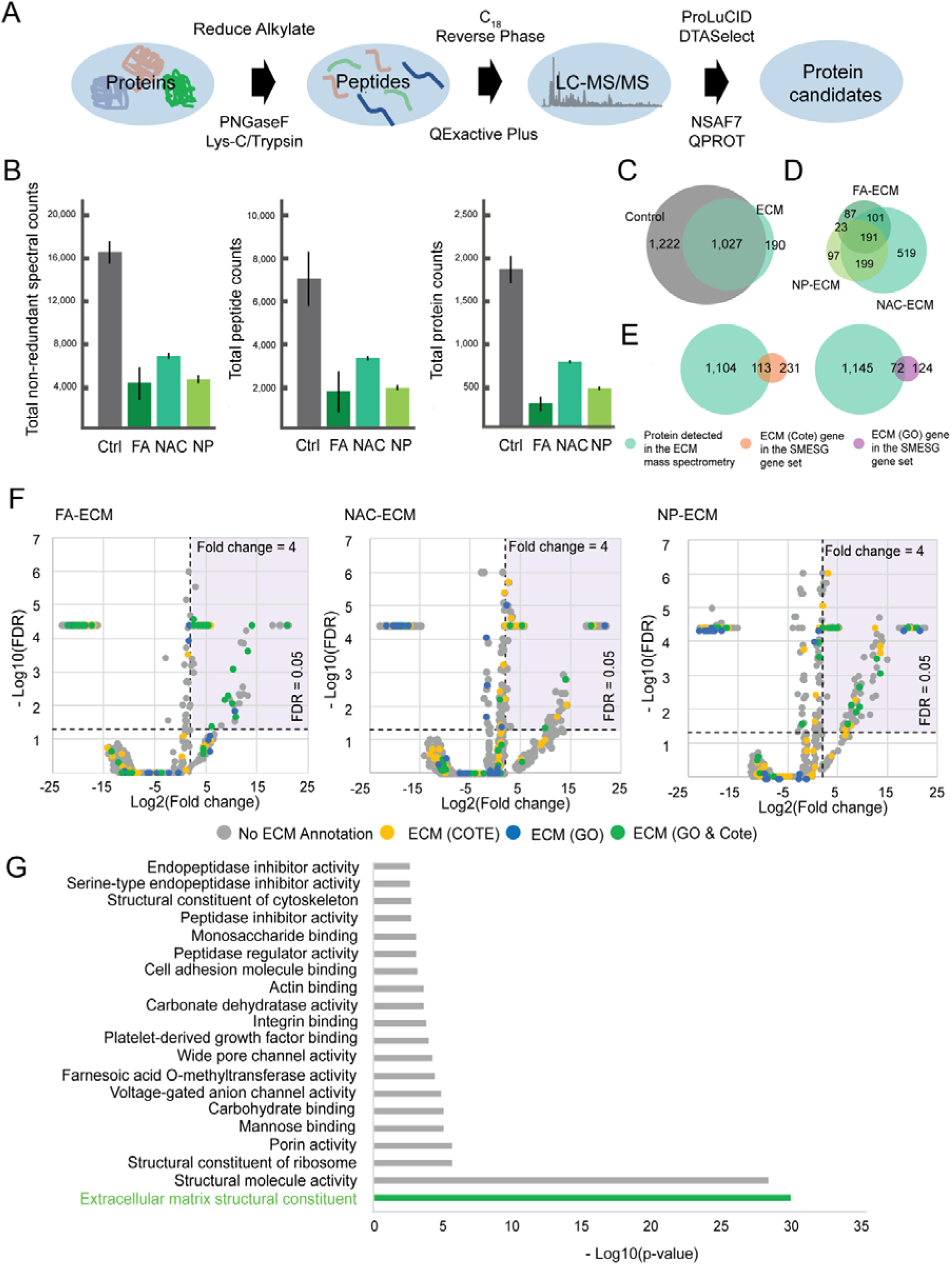
Proteomics analysis reveals a complex and enriched ECM profile from decellularized planarians. (A) Workflow for ECM bottom-up proteomics analysis spectrometry (B) Histograms of the non-redundant spectral, peptide, and protein counts identified in control (Ctrl) and FA-ECM (FA), NAC-ECM (NAC), and NP-ECM (NP) samples. Control, FA-ECM, and NAC-ECM were analyzed in triplicates while NP-ECM had 2 replicates. Error bars represent standard deviation. (C) Venn diagram displays the overlapping proteins identified from ECMs and control, (P < 0.01) (D) Venn diagram represents the overlapping proteins identified from three different ECM isolation procedures. The P value between FA-ECM vs NAC-ECM, FA-ECM vs NP-ECM, and NAC-ECM vs NP-ECM is less than 0.01. (E) Venn diagram displays the overlap between every protein identified from mass spectrometry and ECM identified by Cote et al. (ECM(Cote)) and ECM identified in our work (ECM(GO)) (P < 0.01) (F) Volcano significance plots of the log_2_ fold-change and −log_10_ false discovery rate (FDR) calculated by QPROT for the pairwise comparisons of each ECM from different decellularization procedures compared to the control. The grey dots represent proteins that have no ECM annotation; the yellow dots represent proteins that have been annotated as ECM by Cote et al., while the blue dots represent proteins that have been annotated as ECM by our annotation; the green dots represent proteins that have been annotated as ECM both by Cote et al. and our annotation. Dashed lines are shown at fold change equal to 4 and FDR equal to 0.05. Proteins in the upper right quadrant (purple area) were significantly enriched in the ECM samples versus the control (n=220). The dots were colored based on the annotation. (G) Bar graph represents the value of −log_10_(p-value) of each gene ontology (molecular function, MF) identified in enriched ECM sample.

The mass spectrometry data identified 2,249 proteins in the control sample while combining all three types of ECM samples led to 1,217 identified proteins (Figure 2C). Approximately 1,027 proteins were found in both control and ECM while only 190 proteins were exclusively found in the ECM samples, and 1,222 were found exclusively in the control samples. Some notable proteins found in the ECM samples are cathepsin B (SMESG000048413.1), collagen (SMESG000016282.1), hemicentin (SMESG000021248.1), matrix metalloproteinases (SMESG000049445.1), and nidogen (SMESG000016993.1). The full list of proteins identified in each condition is provided in Table S2. This discovery suggests that our protocol is able to prevent protein degradation and is compatible with mass spectrometry applications.

Since the morphology of isolated ECMs varied depending on the isolation protocol (Figure 1B-D), we tested whether the different ECM isolation protocols could result in differential identification of peptides in our mass spectrometry data (Figure 2D). There was a large overlap of identified proteins between the NAC-ECM and NP-ECM samples. The second largest overlap was seen between the sample of NAC-ECM and FA-ECM, while the least overlap was between FA-ECM and NP-ECM. More importantly, 191 proteins were shared between all three ECM isolation methods while 87, 519, and 97 proteins were exclusively detected in FA-ECM, NAC-ECM, and NP-ECM, respectively. Hypergeometric tests confirm that the overlaps between conditions is greater than expected by chance (p < 0.01). In summary, our planarian decellularization protocol allows us to identify probable ECM proteins and to construct the first experimentally-based matrisome database for planarians.

### Published ECM annotations validate our proposed ECM protein database

In order to evaluate the quality of our data, we compared our mass spectrometry enrichment results with two sets of predicted planarian ECM genes. The first set is a previously published ECM gene set, ECM(Cote) (Cote, Simental, and Reddien 2019). The second predicted ECM gene set is called “ECM(GO)”, and was generated by identifying genes we were able to annotate from the Gene Ontology term “Extracellular Matrix” or any of its child terms. We find a significant degree of overlap between the two datasets (Figure S2) (hypergeometric p < 0.01). This analysis suggests that the two datasets are highly similar.

The set of proteins detected in the mass spectrometry dataset shares 113 members with ECM(Cote) and 72 members with ECM(GO) (p < 0.01 for each) (Figure 2E). This suggested that the mass spectrometry results are consistent with the two bioinformatic predictions. Moreover, we see that many of the genes bioinformatically predicted as ECM do not meet our FDR threshold. This suggests that while we are clearly able to enrich for ECM, we likely have not reached saturation. As more spectra were captured, more ECM genes were discovered in our ECM sample. This orthogonal validation with independent datasets supports our conclusion that we have successfully identified and enriched for ECM components.

### Confirmation of ECM protein enrichment in decellularized samples

The mass spectrometry results show that many ECM proteins were enriched over controls, such as different types of collagen protein (SMESG000066546.1), mucin (SMESG000041532.1), and nidogen 2 (SMESG000016992.1) (Figure 2F and Table S2). Nonetheless, several proteins annotated as ECM components were not significantly enriched, such as matrix metalloproteinase 1 (SMESG000040780.1) and matrix metalloproteinase 2 (SMESG000014326.1). Interestingly, the proteins significantly decreased in ECM samples include lectin 2c (SMESG000056364.1) and mannose-binding protein C (SMESG000046618.1). These observations suggest that although we were able to enrich for ECM proteins, some ECM proteins are not enriched and some are lost during tissue decellularization.

In order to verify the enrichment of ECM proteins in our samples, gene ontology (GO) analysis was performed for proteome data of all the ECM samples. The results further confirmed that the GO term with the highest −log_10_(p-value) was ‘extracellular matrix structural constituent’ (Figure 2G), indicating that these molecules are the major class of proteins in our samples.

We took this opportunity to confirm whether our decellularization method could eliminate intracellular proteins in addition to enriching ECM proteins. Although we experimentally verified the depletion of intracellular proteins as shown by Western blot (Figure 1G), we performed bioinformatic verification by inspecting the statistical distribution of ribosomal proteins, highlighted in red on the volcano plot from FA-ECM samples (Figure S3). We found that 61 out of the 62 detected ribosomal proteins were located outside the upper-right quadrant (Fisher’s Exact p < 0.01), indicating that the majority of ribosomal proteins were not enriched in ECM compared to control. This result highlights the potential of our decellularization procedure to deplete ribosomal proteins and, presumably, other intracellular proteins.

In conclusion, although our ECM proteomic inventory is likely not complete due to the unavoidable loss of certain proteins during sample preparation, we anticipate that our mass spectrometry data is of high quality as our decellularization process does not merely eliminate intracellular proteins, but enriches for *bona fide* ECM proteins as well.

### RNAi screen of planarian ECM identified from proteomics uncovers two genes crucial for tissue homeostasis and regeneration

The proteomics of decellularized planarians offered the possibility of initiating an RNAi screen to discover novel ECM genes responsible for tissue homeostasis and regeneration in planarians (Figure 3A). Forty five proteins were selected from our proteomic inventory for the RNAi screen as well as one previously published control ECM protein (tolloid-like protein 1) that was not present in our inventory (Reddien et al. 2007). The list of all genes targeted in the RNAi screen are listed in Table S3. The worms were fed with bacteria expressing dsRNA of the target gene every three days for a total of ten feedings (Figure 3B). Upon completion of the RNAi feeding schedule, planarians were subjected to two assays to examine the role of the genes of interest during homeostasis and regeneration (Figure 3C).

**Figure 3.**
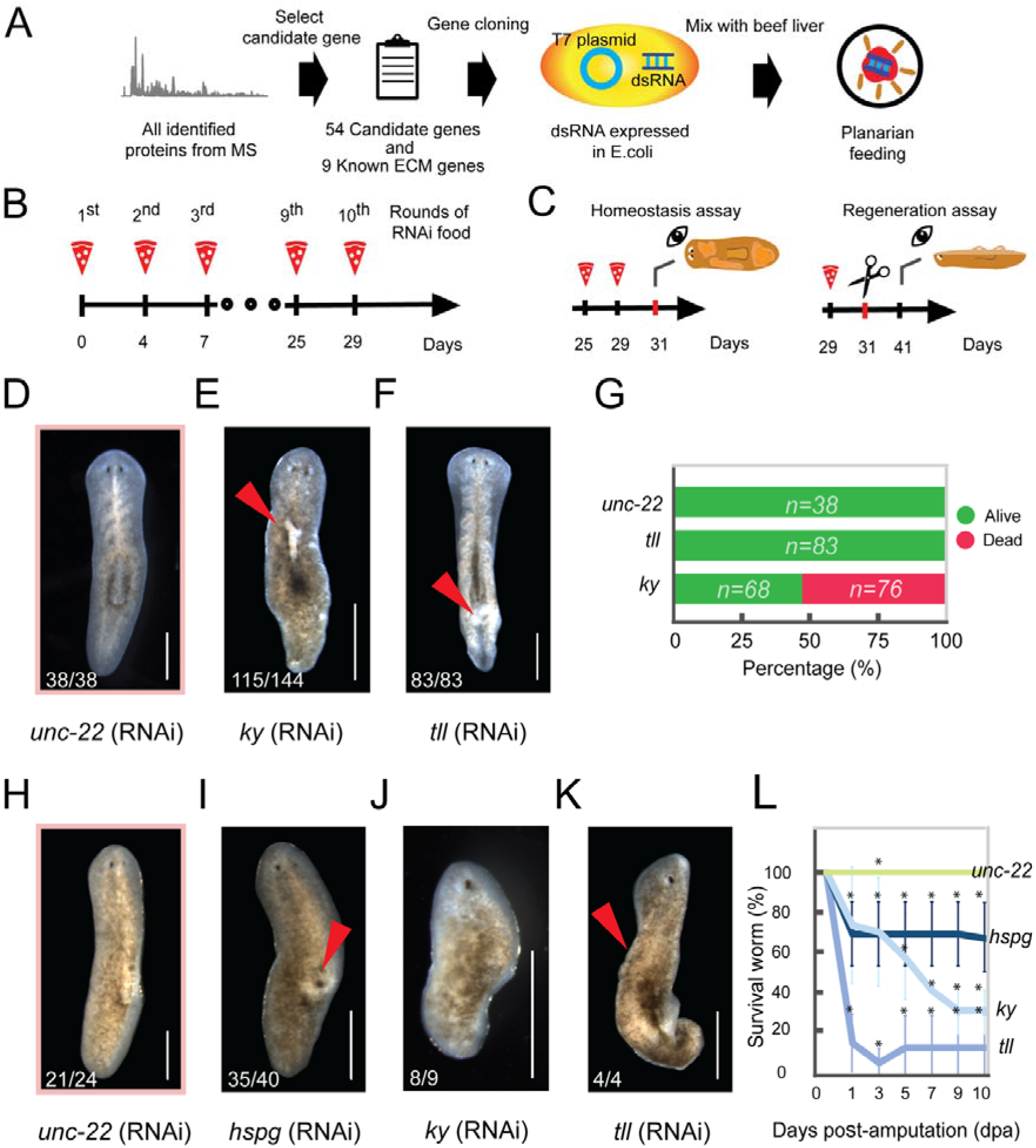
RNAi screening of empirical matrisome discovers two genes of interest involved in tissue homeostasis and/or regeneration. (A) Workflow for dsRNA feeding for RNAi screening. (B) dsRNA feeding schedule. (C) Cartoon illustrating the timeline for homeostasis and regeneration assays. (D-F) Characteristic phenotypes of planarians in homeostasis assay. Scale bars represent 500 μm. (D) Phenotype of planarians with RNAi against *unc-22* (RNAi) as negative control. (E) Phenotype of planarians with RNAi against kyphoscoliosis peptidase (*ky*). The arrow indicates tissue lesion. (F) Phenotype of planarians with RNAi against tolloid-like protein 1(*tll*). The arrow indicates tail curling with rough surface. (G) The survival rate of unamputated planarians by the end of RNAi feeding course. (H-K) Characteristic phenotypes of planarians in regeneration assay. Scale bars represent 500 μm. (H) Phenotype of planarians with RNAi against *unc-22* (RNAi) as negative control. (I) Phenotype of planarians with RNAi against basement membrane heparan sulfate proteoglycan (*hspg)*. The arrow indicates the hump-like structure. (J) Phenotype of planarians with RNAi against kyphoscoliosis peptidase (*ky*). (K) Phenotype of planarians with RNAi against tolloid-like protein 1(*tll*). The arrow indicates rough dorsal surface. (L) Survival rate of RNAi-fed planarians 1, 3, 5, 7, 9, and 10 days after sagittal amputation. Error bars represent standard deviation. Stars indicate statistically significant differences, as calculated by ANOVA (* = p < 0.05), compared to *unc-22* as the negative control. Biological replicates of RNAi-treated worms of *unc-22*, *hspg*, *tll*, and *ky* worms were 3, 6, 5, and 3 respectively.

Our RNAi screen of 45 genes identified from mass spectrometry found that knockdown of one gene encoding kyphoscoliosis peptidase (*ky*) caused visible phenotypes in animals at homeostasis comparing to negative control. Body lesions were observed on the dorsal surface as indicated with the arrow (Figure 3D and E). In addition, our RNAi knockdown of the gene encoding tolloid-like protein 1 (*tll*) also caused abnormal phenotypes in animals at homeostasis. The tails of *tll*-knockdown worms appeared to be rough, as indicated by the arrow (Figure 3F). Moreover, there were three parallel dark streaks present on the dorsal surface of the animal (Video S2). The animals curl along the outer two streaks resulting in the observed tube-like phenotype at the posterior. We scored for worm death at the end of RNAi feedings and found that 53% of *ky*-knockdown worms died while *tll*-knockdown worms did not. (Figure 3G).

After 10 RNAi feedings the animals were amputated sagittally to score for regeneration. This screen identified roles for two genes encoding the basement membrane heparan sulfate proteoglycan (*hspg*) and kyphoscoliosis peptidase (*ky*) (Figure 3H-J). Negative controls, *unc-22* RNAi animals, showed normally regenerated tissue, as indicated by the presence of eye spots on the new tissue with an unpigmented blastema (Figure 3H). In contrast, the *hspg*-knockdown worms showed a bulging structure at the pharyngeal position, indicated with the arrow, while the regenerated blastema and eye spots were normal. (Figure 3I). Secondly, although *ky*-knockdown worms were able to form new blastemal tissue, the eye spots were not visible (Figure 3J). As expected, *tll*, which previously showed a phenotype in the homeostasis assay, was also found to have a phenotype in the regeneration assay. The worms failed to produce a blastema and eye spots, indicated by a lack of unpigmented tissue at the wounded edge; the dorsal surface of the planarians also appeared to be rough, indicated with a yellow arrow (Figure 3K). Moreover, the *tll(RNAi)* animals were ventrally curled, particularly at the tail region, in a manner that was similar to the analogous worms at homeostasis (Figure 3F). In total, three ECM genes of interest were identified from both assays and the reduction of gene expression in all RNAi conditions was confirmed by quantitative polymerase chain reaction (Figure S4).

Because some animals died during the regeneration assay, we then conducted a survival assay to quantify the lethality of these three RNAi conditions. All three conditions showed reduced survival during regeneration, while 100% of the control animals fully regenerated and survived (Figure 3L). Only 15% of *tll*-knockdown worms survived at 1 day post-amputation (dpa), while the percentage of survival for *hspg*-knockdown worms was 68% at the same time point. In contrast, the survival of *ky*-knockdown worms gradually decreased from 1-dpa to 10-dpa. At the endpoint, the survival of RNAi-treated worms at 10-dpa were 67%, 10%, and 30% for *hspg*, *tll*, and *ky*, respectively. These results indicated that the knockdown worms of all three genes of interest are more sensitive to injury than control animals since their survival rates dropped sharply at the first day after amputation. This suggests that these genes could be required for the survival of planarians at the early stage of tissue injury.

To summarize, we conclude that *ky* is essential for tissue homeostasis and regeneration while *hspg* is important for tissue regeneration. Because we were able to replicate the previously reported phenotype for the predicted ECM gene *tll*, we conclude that our RNAi screening is reliable and could provide insights on contribution of a particular ECM genes in homeostasis and regeneration.

### Analysis of protein sequences reveals structural and functional domains

In order to further investigate the function of the genes of interest, the protein domain composition of all three proteins was bioinformatically predicted (Figure S5). The domain analysis showed that the HSPG protein contained several domains such as immunoglobulin (IG), low-density lipoprotein receptor domain class A (LDLa), laminin B domain, and epidermal growth factor-like domain (EGF). Secondly, the KY protein only contains two repeated domains of transglutaminase/protease-like homologues (TGc). Lastly, the TLL protein possesses a zinc-dependent metalloprotease domain with several repeats of CUB domains and calcium-binding EGF-like domain (EGF-Ca). These results suggest the potential function of each protein behind our discovered phenotypes in knockdown animals: *hspg* likely encodes a large core matrisome structural protein, while the protein products of *tll* and *ky* are likely matrisome-associated metalloproteases.

### Expression patterns of genes of interest

Because we discovered roles for ECM genes in regeneration, we therefore sought to determine whether these genes were expressed in stem cells. We used γ-irradiation to eliminate neoblasts, followed by whole mount *in situ* hybridization to visualize the gene expression patterns (Figure 4A). Prior to irradiation, *hspg* is expressed all over the body of planarians, especially at the pharynx. *ky*, however, was found to be expressed prominently in the planarian digestive tract. The expression of *tll* was also found to be well distributed in every part of the worm body. Interestingly, while irradiation eliminated the probe signal for our neoblast marker, *piwi-*1, it did not eradicate the probe signals for any of the genes of interest, suggesting that they are not expressed in stem cells.

**Figure 4.**
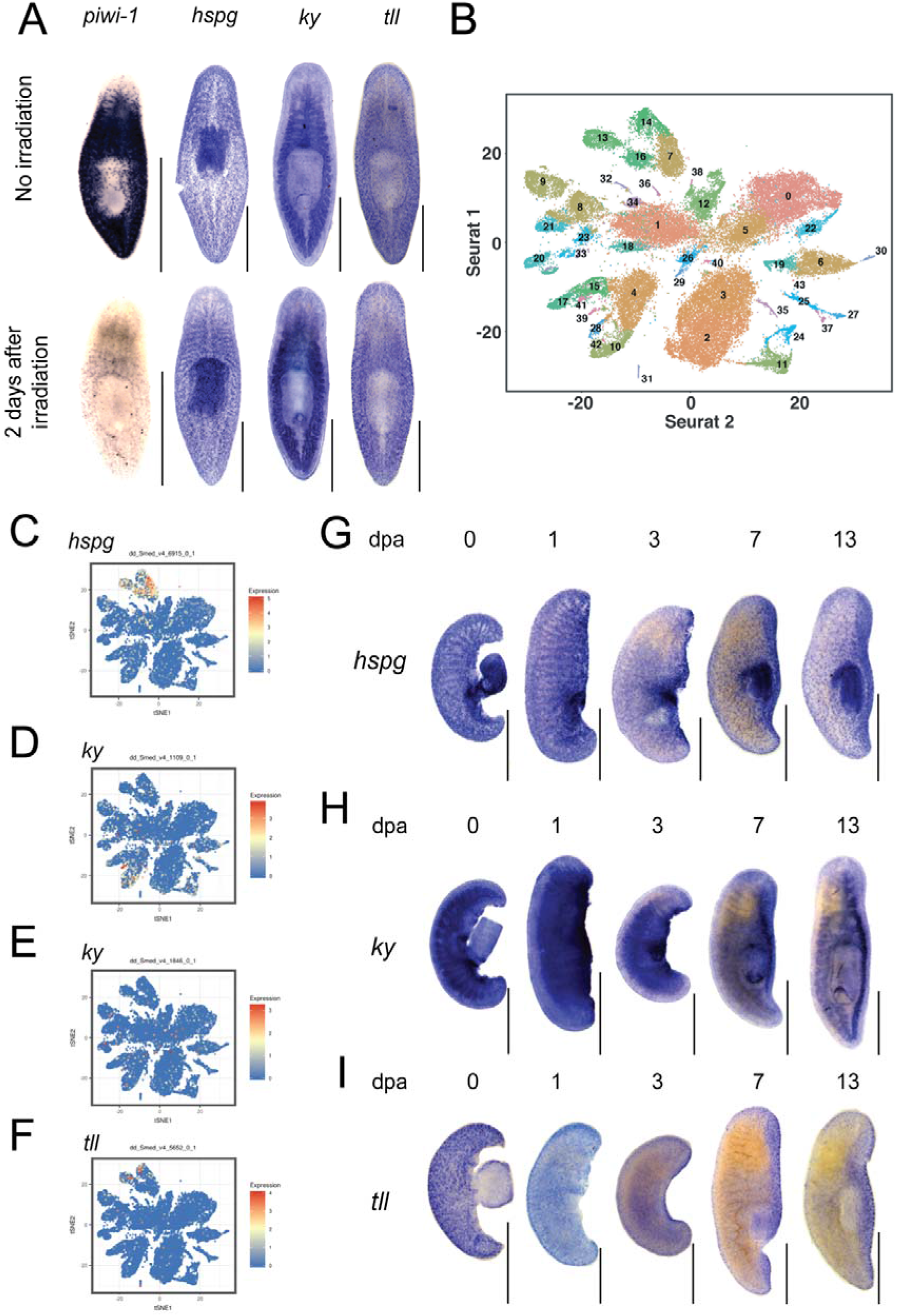
Spatiotemporal characterization of genes of interest during tissue homeostasis and regeneration. (A) *In situ* hybridization of genes of interest on unamputated planarian prior to, and two days after, irradiation. Scale bars represent 500 μm. (B) The reference atlas of RNA single-cell sequencing published by Fincher and colleagues (Fincher et al. 2018). (C-F) The relative expression of (C) *hspg*, (D and E) *ky*, and (F) *tll* in each cell type in the RNA single-cell sequencing. Different dot color indicates differences in level of gene expression. (G-I) *In situ* hybridization of (G) *hspg*, (H) *ky*, and (I) *tll* on sagittally amputated planarians regenerated for 1, 3, 7, and 13 days. All scale bars represent 500 μm.

In order to pinpoint the cell type responsible for expressing each of our genes of interest, we consulted the planarian single cell RNA sequencing database *Digiworm* (Fincher et al. 2018) (Figure 4B). The results showed that *hspg* and *tll* were highly expressed in muscle cells, while *ky* was enriched in many cell types such as cathepsin+ cells, epidermis, and intestine (Figure 4C-F). Both WISH and single cell RNA sequencing suggest that all three genes of interest are mainly expressed in differentiated cells such as gut, muscle, or epidermis.

As all three genes of interest affect tissue regeneration in amputated planarians, we used *in situ* hybridization to determine if the expression patterns of these genes change during regeneration (Figure 4G-I). While the original pattern of *hspg* remained, after 1-dpa there was an increase in expression at the location of the original pharynx (Figure 4G). At 3-dpa, *hspg* global expression was reduced, while prominent expression of *hspg* remained in the newly formed pharynx area. The signal of *hspg* expression in the pharynx was persistent throughout the regeneration process. Additionally, *ky* was strongly expressed in the newly regenerated blastema, especially in the intestine near the wounded area at 1 and 3-dpa. At 7- and 13-dpa, *ky* expression was reestablished at the newly regenerated blastema particularly in the area that forms the regenerating intestine (Figure 4H). Lastly, the expression of *tll* decreased globally at 1-dpa and only a slight signal of *tll* expression was observed at the edge of the wound (Figure 4I). However, the signal reemerged in the newly regenerated blastema at 3-dpa and its expression was persistent particularly at the newly regenerated blastema from 7-dpa onward. Our results suggest that the three genes are expressed mostly in differentiated cells such as muscle or epithelial cells. Moreover, all of our genes of interest respond to tissue amputation as shown by dynamic changes in their gene expression patterns following wounding. Taken together, these results indicate the role of the three genes of interest in the physiological response of planarian regeneration.

Ultimately, we have demonstrated the potential of our newly-described decellularization procedures for isolating high-quality ECM from planarians for use in many downstream applications. Here, we showed the isolated ECM can be used for a protein mass spectrometry pipeline, from which the first experimentally-based ECM proteomic database has been established. This work empowers systematic study of planarian ECM to characterize its function in tissue homeostasis and regeneration.

## Discussion

The study of ECM in a highly regenerative animal model, such as planarians (Reddien and Sánchez Alvarado 2004), is currently limited, preventing us from understanding how the ECM may or may not contribute to tissue regeneration (Isolani et al. 2013; Seebeck et al. 2017; Lindsay-Mosher, Chan, and Pearson 2020). To resolve this limitation, we established an integrative workflow consisting of tissue decellularization, proteomics, and RNA mediated genetic interference (RNAi) for the purification and characterization of planarian ECM.

The quality of an isolated ECM is a significant parameter for its downstream characterization. Our decellularization procedure resulted in the production of a full-body ECM scaffold from *S. mediterranea* possessing both little residual genomic DNA and a well-defined basement membrane (Figure 1). Both of these parameters are generally used as gauges for determining the quality of purified ECM after intensive decellularization (Gilbert et al. 2009; Hynes 2009; Ross et al. 2009; Nakayama et al. 2010). Not only does our procedure produce an isolated ECM of high quality, but it also suggests that a single protocol with a few customizations may ultimately be applicable to a broad diversity of planarians (Sonpho et al. 2020).

Mass spectrometry of the purified ECM allowed us to establish an experimentally-defined planarian matrisome inventory. When compared to a computationally-derived ECM catalogue (Cote, Simental, and Reddien 2019), we found both an overlap of 113 proteins between databases, with some proteins represented in our matrisome but not in the *in silico* inventory 1,104 proteins and vice versa 231 proteins (Figure 2E). For example, we found lectin 1 (SMESG000028263.1), cathepsin L1-like (SMESG000049722.1), and laminin subunit β (SMESG000062589.1) as being shared, but found prosaposin (SMESG000019976.1), and Fucolectin-4 (SMESG000059878.1) missing in the computationally-derived matrisome inventory and some well-defined ECM proteins, such as lysyl oxidase 3, netrin 5, and fibrillin 2, missing from the experimentally-defined matrisome. There are several likely explanations for these discrepancies. First, our decellularization protocols are not only effective for removing cells but also for enriching ECM and ECM associated proteins (Figure 2F and G) that are not likely to be pre
dicted computationally. Second, mass spectrometry is likely to be unable to reliably detect proteins that may only be present in the sample in small amounts, with their signals overwhelmed by the overabundance of major ECM proteins such as collagen (Lampi et al. 2005), which may be addressed experimentally in the future by collagenase treatment prior to analysis. Third, there may be unintended loss of ECM proteins in the preparation process, in particular during the extensive washing steps (Li et al. 2016). And a fourth possibility for the discrepancies observed may the presence of unannotated genes in *S. mediterranea* (Rozanski et al. 2019) indicated in Table S5, suggesting that our ECM characterization workflow stands to uncover novel, uncharacterized components of the planarian ECM. A full characterization of those unknown proteins is beyond the scope of this study. Our research group is looking forward to characterizing those unknown proteins in a future work.

Among the three decellularization schemes, our data also indicated that the different ECM preparations may affect the quality recovery of proteins (Figure 2D). For instance, the FA-ECM dataset had the lowest protein count, possibly due to the intensive formaldehyde crosslinking of the sample, which likely resulted in the exclusion of large cross-linked protein complexes from MS analysis. The differences of proteomic profiles and the number of identified proteins observed among the different methods employed for decellularization are in good agreement with previously published analyses (Didangelos et al. 2010; Coronado et al. 2017). Hence, our findings indicate that the selection of a given decellularization method over another must be guided by the downstream applications for the purified ECM.

Once we were able to experimentally define a planarian matrisome inventory, we sought to test the role of these components in both tissue homeostasis and regeneration (Figure 3). To our knowledge, the genes *hspg* and *ky* have not yet been characterized in planarians. Our known ECM protein, *tll*, has only been previously reported to influence the formation of tissues at the midline and pattern tissues along the dorso-ventral axis (Reddien et al. 2007) and predicted to be an ECM protein (Cote, Simental, and Reddien 2019). Previous work indicates that the muscle is responsible for most of the ECM production and secretion (Cote, Simental, and Reddien 2019), yet there is little information available about the distribution of secreted ECM components across the planarian body. Whole-mount *in situ* hybridization and single cell RNA sequencing database cross-referencing support that the products of the *hspg*, *tll* and *ky* genes are expressed in differentiated cells (Figure 4A-F). In addition, our data indicated that expression of all three genes of interest is both quantitatively and locally dynamic during regeneration (Figure 4G-I), similar to the behavior of previously studied ECM genes (Wurtzel et al. 2015).

The expression of *hspg* during both homeostasis and regeneration is mostly localized to muscle (Figure 4A-C and G), particularly in the pharynx which is mainly composed of muscle cells (Kobayashi et al. 1998). RNAi against *hspg* clearly results in defective regeneration as a humped-like structure is observed where the pharynx should be located (Figure 3I). We therefore hypothesize that *hspg* likely plays a role in pharynx regeneration or that this gene might be involved with regulation of the basal lamina structure since its gene expression is similar to that of *megf6* and *hemicentin,* which are reported to be important for maintaining basement membrane integrity (Lindsay-Mosher, Chan, and Pearson 2020). Secondly, the knockdown of *ky* expression results in the epithelial lesions (Figure 3E), which demonstrate the function of *ky* in epithelial homeostasis or differentiated cells. Additionally, our data suggested a function of *ky* in intestinal regeneration (Figure 4A, B, D, E and H). It has been reported that there are proteins with protease domains that are expressed in the digestive tract and play roles in tissue regeneration (Goupil et al. 2016), inviting the possibility that the similarly expressed proteins play similar roles in planarians. Moreover, *ky* might be a crucial protein for the function of intestinal cells in that the knockdown of *ky* might cause the malfunction of the intestinal tract (Figure 3J), since it has been reported that starvation impedes the rate of regeneration in planarians (Brøndsted 1953). Finally, *tll* has been previously implicated in regulating dorsal midline patterning in regenerating planarians (Reddien et al. 2007). However, the exact mechanism of how *tll* controls stem cell differentiation at the midline is not fully understood in planarians. As *tll* is part of the BMP pathway, which is essential for dorso-ventral polarity regulation throughout the animal kingdom (Tucker, Mintzer, and Mullins 2008), it is possible that this gene may both serve as a secreted ECM-modification protein in addition to participating in the BMP pathway. The BMP-related functions of *tll* may explain the phenotypes observed on the dorsal surface of planarians (Figure 3F and K), but further biochemical characterization will be needed to understand its association with the ECM.

We provide proof-of-concept that our proposed workflow is effective for studying ECM components involved in planarian tissue regeneration and homeostasis (Figure 5). The discovery of two genes with promising functions and obvious knockdown phenotypes supports the validity of our decellularization-based planarian matrisome approach to target gene candidates for RNAi screening. Our work thus sets the stage for investigating the involvement of different ECM components in complex biological processes, offering a unique approach for examining in detail the effects the microenvironment may play in stem cell fate regulation and determination during tissue homeostasis and whole-body regeneration. More importantly, this workflow provides an opportunity to discover novel ECM proteins that fail to be predicted by bioinformatic approaches because of unique features or specificity to planarians. Ultimately, our work highlights the potential of planarians as a model system for the study of ECM remodeling.

**Figure 5.**
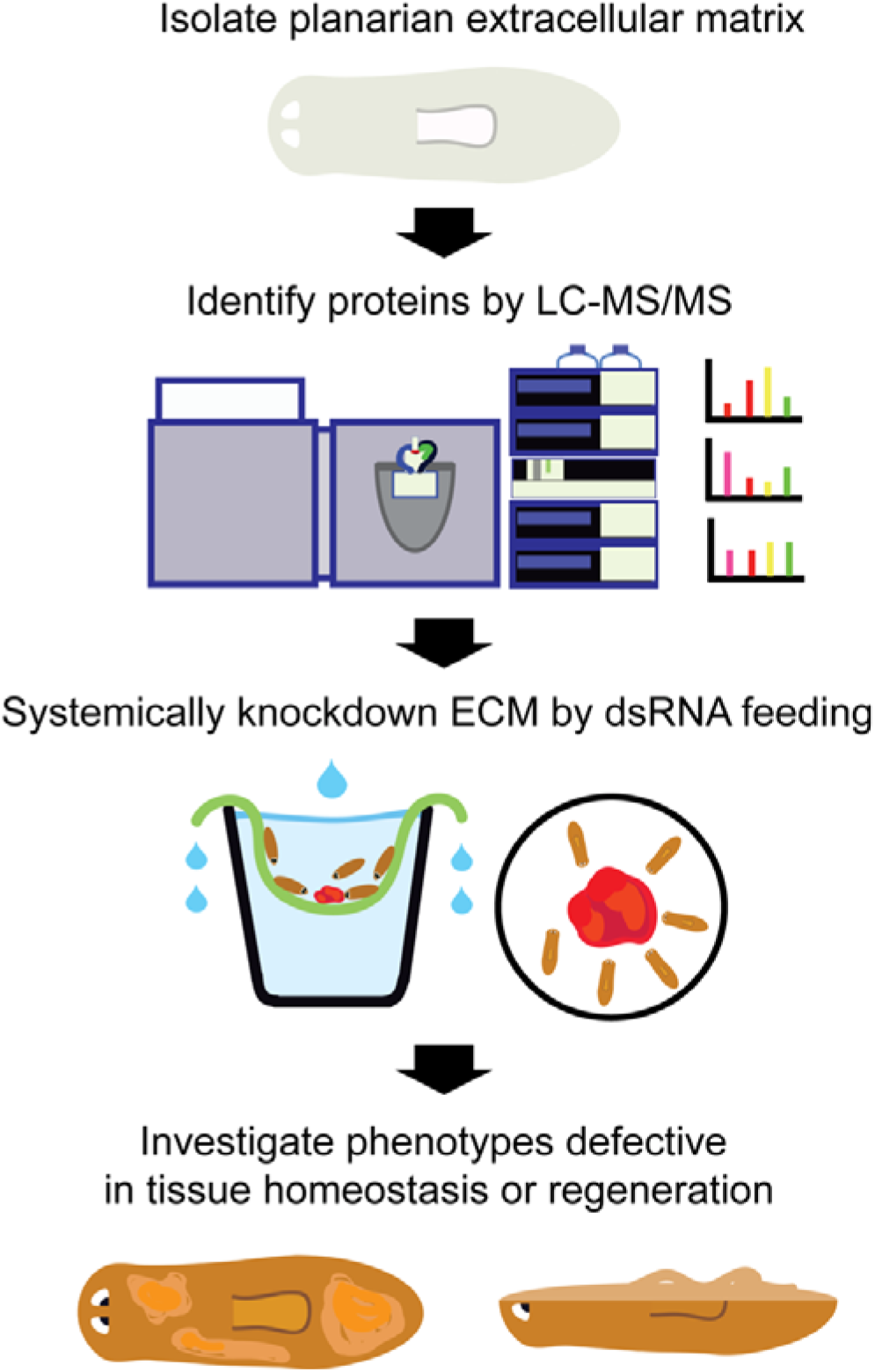
An overview of our pipeline for characterizing planarian ECM to study regeneration, development, and homeostasis. We describe a multidisciplinary workflow to isolate, analyze, and characterize the functions of extracellular matrix components during homeostasis and regeneration in planarians. ECM is enriched through one of three alternative decellularization procedures. The protein composition of the isolated ECM is empirically determined via liquid chromatography coupled to mass spectrometry. The genes encoding these components are perturbed by RNA interference to reveal visible a natomical phenotypes indicating functional roles for the identified ECM proteins in homeostasis or regeneration.

## Acknowledgments

We would like to acknowledge the financial support to Ekasit Sonpho from Junior Research Scholarship Program (JRS), Fulbright Program during his stay at the Stowers Institute for Medical Research, USA. We are grateful for the support received from members of the research core facilities at the Stowers Institute for Medical Research including all planaria core facility personnel; Yan Hao, proteomics center; Nancy Thomas, histology facility; Dan Bradford and Michael Peterson, molecular biology facility; Melainia McClain, electron microscopy center; and Cindy Maddera, microscopy center. We also appreciate constructive comments from Lauren Cote following our preprint publication. Alejandro Sánchez Alvarado, Eric J. Ross and Frederick G. Mann, Jr. are supported by Howard Hughes Medical Institute. We appreciate the valuable input from every ASA lab members. Works in Ounjai lab is supported by the Center of Excellence on Environmental Health and Toxicology (EHT), Faculty of Science, Mahidol Universtiy, and the Thailand Science Research and Innovation (TSRI). Moreover, we also thank Ekarat Sonpho for editing all the video materials, and Mr. Jirawat Salungyu for assistance in statistical analysis.

## Data Availability

The mass spectrometry proteomics data have been deposited to the ProteomeXchange Consortium via Pride (Deutsch et al. 2017; Perez-Riverol et al. 2019) partner repository with the dataset identifier PXD013181 and 10.6019/PXD013181. Original data underlying this manuscript may also be accessed after publication from the Stowers Original Data Repository at http://www.stowers.org/research/publications/libpb-1407.

## Notes

http://www.stowers.org/research/publications/libpb-1407

### Competing Interest Statement

The authors have declared no competing interest.

### Summary of Updates

This new version incorporates comments and suggestions from readers. Also, proteomics and transcriptomic analyses have been updated.

