## Supplemental Figures and legends for "Decellularization enables functional analysis of ECM remodeling in planarian regeneration"

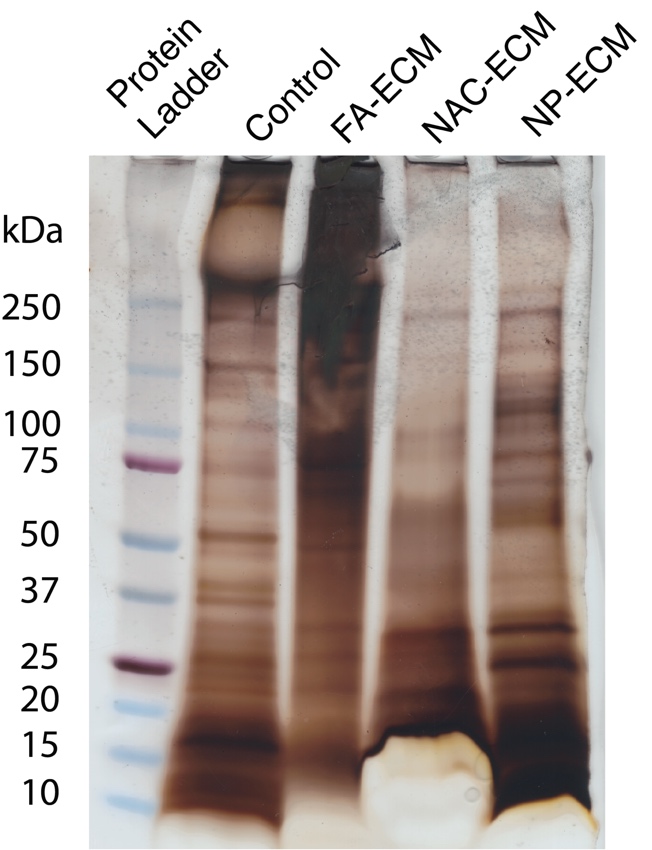

**Figure S1: SDS-PAGE of ECM isolated from *S. mediterranea.*** SDS-PAGE of ECM visualized by silver staining.

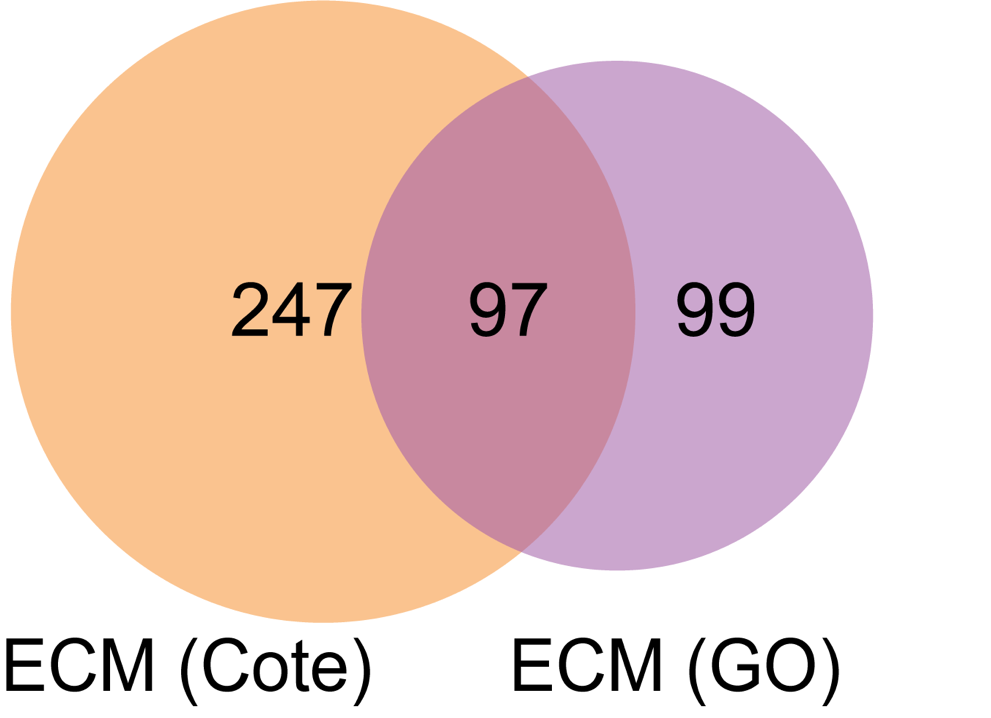

**Figure S2: Venn diagram between two sets of predicted planarian ECM genes.**

ECM (Cote) is a ECM dataset predicted by Cote (Cote, Simental, and Reddien 2019) while ECM (GO) is a ECM dataset predicted in our work (P < 0.01).

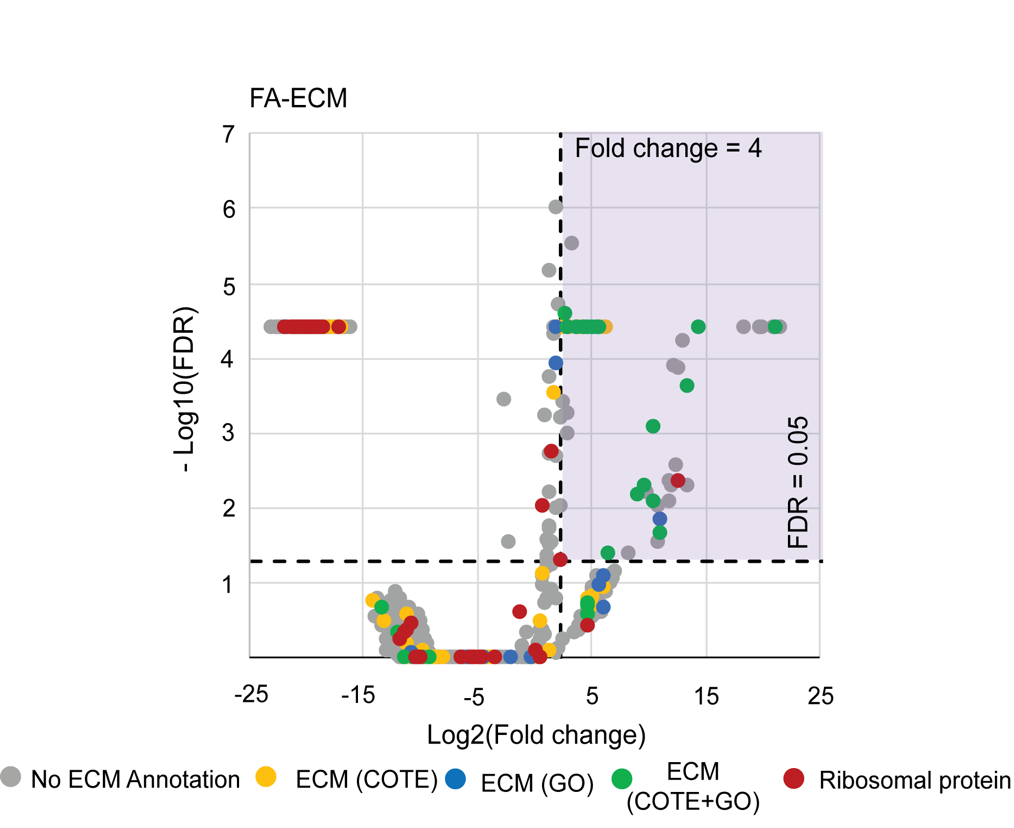

**Figure S3: Volcano plots of FA-ECM compared to control, with highlighted ribosome proteins.** Volcano plots of the log2 fold change and -log10 false discovery rate (FDR) of FA-ECM compared to the control measured by QPROT statistical analysis of FA-ECM compared to control, with 3 triplicate samples for each. The grey dots represent proteins that have no ECM annotation; the yellow dots represent proteins that have been annotated as ECM by Cote et al; the green dots represent proteins that have been annotated as ECM by both our annotation and Cote et al, while the blue dots represent proteins that have been annotated as ECM by our annotation; the red dots represent ribosomal proteins. Dashed lines are shown at log fold change equals to 4 and false discovery rate equals to 0.05. Proteins in the upper right quadrant (purple area) or log_2_(Fold change) ≥ 2 and -log_10_(FDR) ≤ 0.05) were significantly enriched in the ECM samples versus the control (n=220).

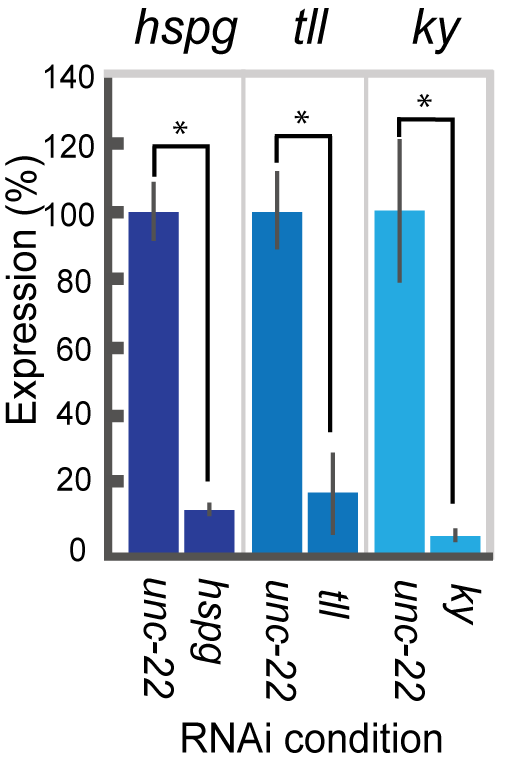
 **Figure S4: RNAi knockdown efficiency.**

The evaluation of RNAi knockdown efficiency of unamputated planarians by qPCR. The worms fed *unc-22* dsRNA were used as negative control for RNAi experiment. Tubulin gene was an internal control for qPCR. Error bars represent standard deviation. The data were calculated from 3 independent experiments, with stars indicating differences statistically-supported by ANOVA (* = P < 0.05).

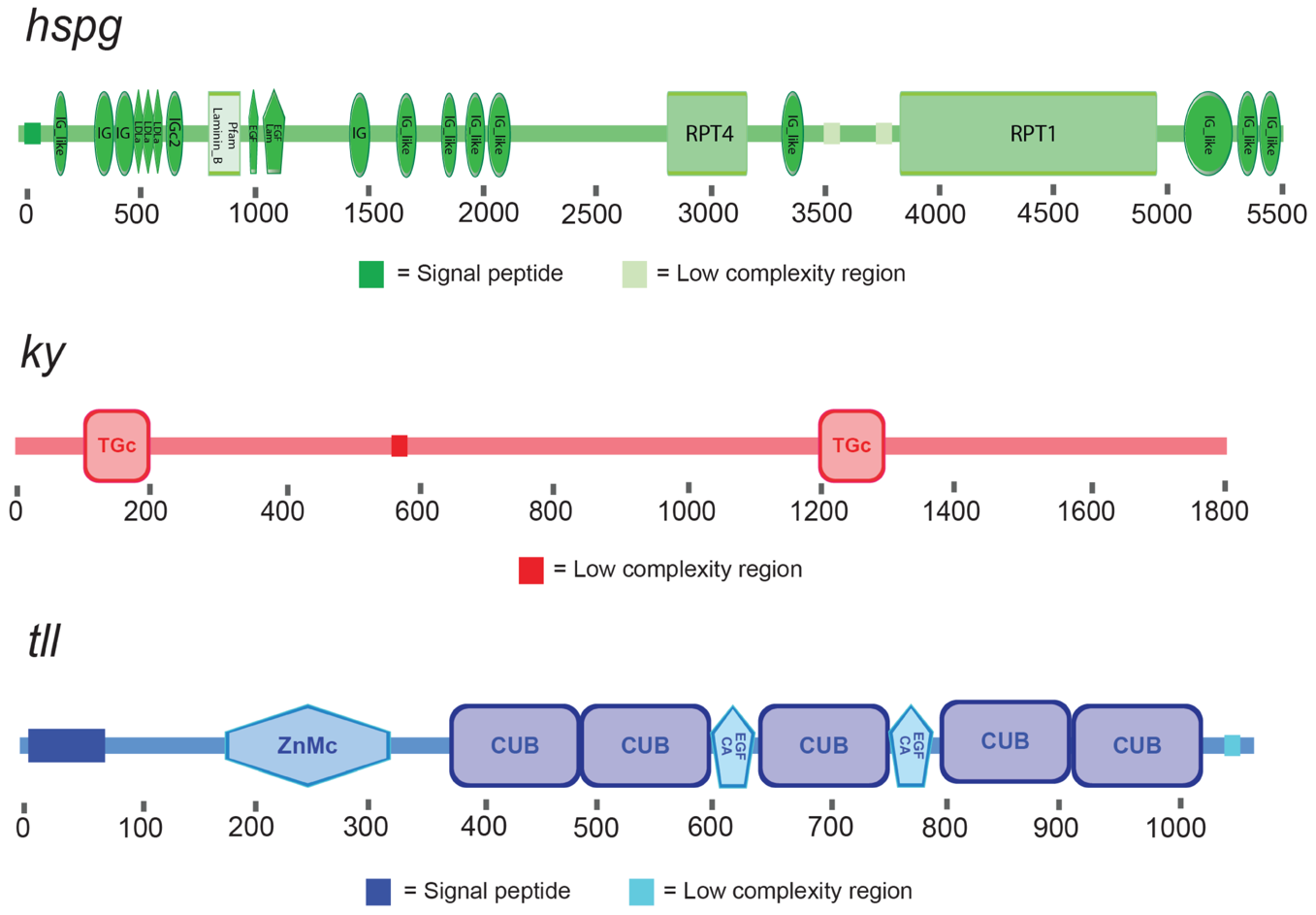

**Figure S4: Domain organization of the three candidate proteins.** Protein domain prediction and assignment of the three selected genes; *hspg*, *tll* and *ky*. The numbers below each line indicate the positions of amino acid in the protein sequence.

**Video S1: Time-lapse of planarian decellularizing by using three different protocols**

The video displays a sped up time-lapse of decellularizing planarians from different decellularization protocols including formaldehyde-treated ECM (FA-ECM), N-Acetyl cysteine-treated ECM (NAC-ECM) and no pre-treatment ECM (NP-ECM). Scale bars represent 1 mm.

**Video S2: Footage of *tll*-knockdown planarians**

The video shows the homeostasis defect of *tll-*knockdown planarians. This worm has been recorded two days after the completion of tenth RNAi feeding. Scale bars represent 1 mm.

**Supplementary Table**

**Table S1:** Chemical compositions of stabilization, decellularization, and wash solutions used to isolate ECM from Schmidtea mediterranea planarians.

**Table S2:** Contrast report and QPROT analysis of the proteins identified in the whole animal (Ctrl), FA-ECM, NAC-ECM, and NP-ECM samples with significant enrichment in the ECM fractions.

**Table S3:** 46 genes candidates for RNAi screening.

**Table S4:** Summary of non-redundant (NR) spectra, peptides and proteins for each experimental replicate analyzed on the QExactive Plus mass spectrometer.

**Table S5:** The number of gene models unannotated (described as Unknown Proteins) and gene models without gene ontology (GO) are shown for each of the datasets. The ECM sample contains the gene models from the FA-, NAC-, and NP-ECM samples combined.
